# Magnetic-alignment of polymer macro-nanodiscs enable residual dipolar couplings based high-resolution structural studies by NMR

**DOI:** 10.1101/671982

**Authors:** Thirupathi Ravula, Ayyalusamy Ramamoorthy

## Abstract

Residual dipolar couplings (RDCs) have been shown to be valuable for the structural studies of systems ranging from small molecules to large proteins. Here we demonstrate the lyotropic liquid crystal behavior of polymer macro-nanodiscs (> 20 nm in diameter) and enable the measurement of RDCs using high resolution NMR.

---

Polymer nanodiscs are being successfully used in structural and functional studies of membrane proteins.^1–6^ A polymer nanodisc is composed of a planar lipid bilayer surrounded by an amphiphilic polymer belt.^7^ A variety of polymers have been used to form nanodiscs using synthetic lipids or by directly extracting membrane proteins with their native lipid environment.^2, 8–9^ Starting with styrene maleic acid,^10^ there has been several different polymers^11–13^ and their derivatives^14–19^ that have been used to overcome the limitations such as size control, metal ion and low pH instability.^14–15, 20^ Furthermore the size control enabled the use of macro-nanodiscs (> 20 nm) in solid-state NMR due to their magnetic alignment properties in the presence of an external magnetic field.^14–15, 21^ In this study, we exploit the magnetic-alignment property of macro-nanodiscs for high-resolution NMR studies of water-soluble proteins using residual dipolar couplings (RDCs).

One of the powerful techniques used in the structural characterization of biomolecules is RDCs. RDCs have been shown to provide valuable global orientation constraints to determine and refine high-resolution structures of biomolecules.^22–24^ RDCs are typically measured using an anisotropic environment created by magnetically oriented liquid crystalline molecules or mechanically compressed/stretched gels.^23, 25^ Previous studies have shown that the degeneracy of bond orientations obtained from RDC values can be minimized using orthogonal tensors by measuring RDCs in different alignment media or modulating the interaction between the alignment medium and protein (or molecular system under investigation).^26^ Bicelles have been used to achieve the measurements of orthogonal alignment tensors by changing the net charge via doping with negatively charged lipid^26^ or surfactants.^27^ However, bicelles are not usable for many biomolecular systems due to the presence of a denaturing detergent. In this study we show that polymer nanodiscs can be used as an alignment medium for measuring RDCs. Our results demonstrate that polymer nanodiscs exhibit a highly ordered homogeneous alignment in the presence of a magnetic field, and the degree of alignment can be scaled by the concentration as well as temperature of the sample. Using cytochrome c (cyt c) as a model system, we show that the experimentally measured RDCs are in agreement with values calculated from reported structures.

Styrene maleic acid quaternary ammonium (SMA-QA) polymer was synthesized as described previously.^15^ Polymer macro-nanodiscs were prepared using 1:0.5 w/w ratio of DMPC and SMA-QA (Figure 1A). The resulting nanodiscs were purified using size exclusion chromatography (SEC), and characterized by dynamic light scattering (Supporting information Figures S1 and S2) and transmission electron microscopy (TEM). The TEM images showed the presence of macro-nanodiscs that were ~25±4 nm in diameter (Figures 1B and S3).

**Figure 1.**
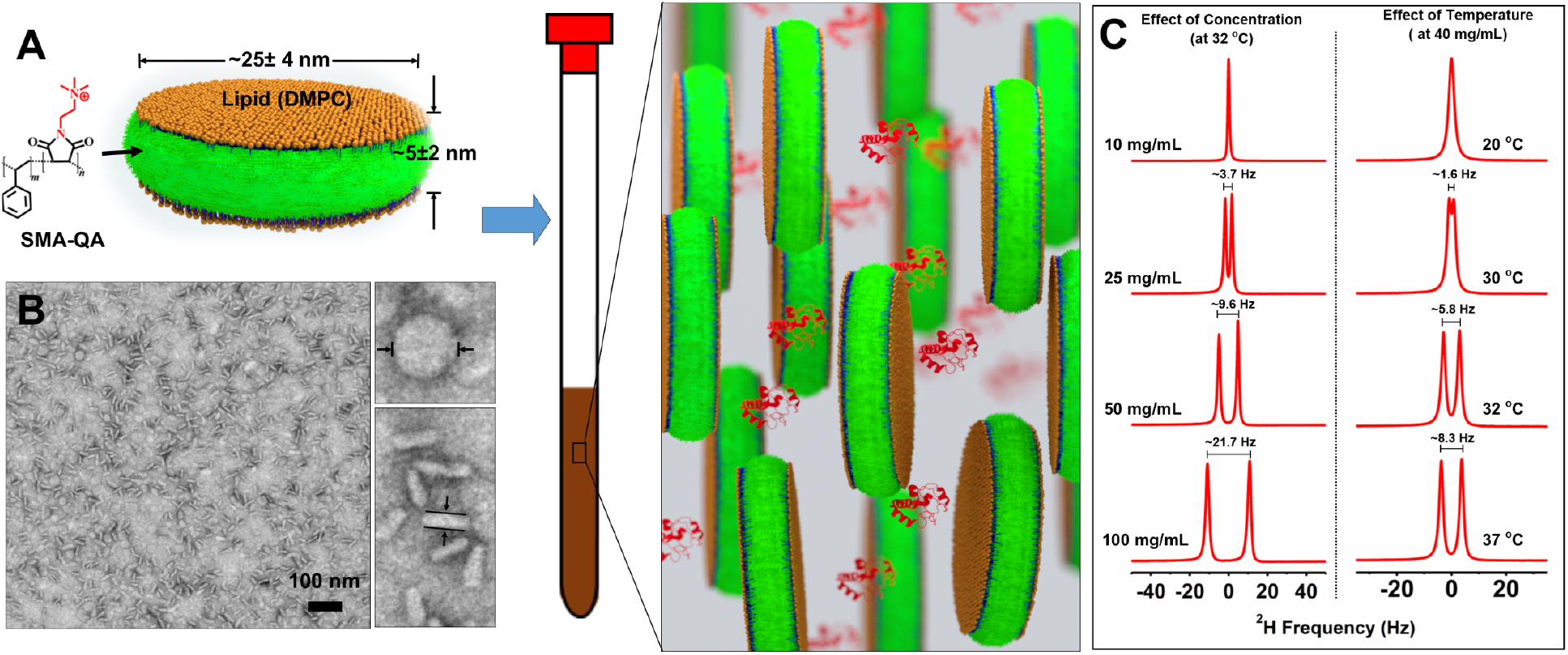
A) Schematic of a polymer lipid bilayer nanodisc and a representation of its use as an alignment medium. B) Transmission electron microscope image of macro-nanodiscs prepared using 1:0.5 w/w ratio of DMPC:SMA-QA. C) Deuterium NMR spectra of macro-nanodiscs with a varying lipid concentration (at 32 °C) and at different temperatures (at 40 mg/mL). All spectra were acquired using a 500 MHz NMR spectrometer.

Macro-nanodiscs with a varying lipid concentration (10-100 mg/mL) were prepared in a 10% D2O containing phosphate buffer. ^2^H NMR was used to examine the anisotropic property of macro-nanodiscs. An isotropic peak was observed at a low lipid concentration (10 mg/mL) at 32° C, whereas by increasing the concentration to 20 mg/mL a doublet was observed with ~3.7 Hz ^2^H quadrupolar splitting (Δν_Q_) from HOD (Figure 1C). Further increase in the lipid concentration to 100 mg/mL increased the observed ^2^H quadrupole splitting to ~21.7 Hz. These observations suggest that the degree of alignment of macro-nanodiscs is scalable by altering the lipid concentration; and the observed symmetric, narrow peaks (line width < 2 Hz) indicate a highly ordered homogeneous alignment of nanodiscs. The effect of temperature on the alignment was examined using 40 mg/mL lipid concentration of nanodiscs. An isotropic peak was observed at 20 °C (which is below the gel-to-liquid crystalline phase transition temperature of DMPC), while a doublet with a quadrupole splitting of ~1.6 Hz appeared as the sample temperature increased to 30 °C. Further increase in temperature increased the observed quadrupole splitting as shown in Figure 1C suggesting that the macro-nanodiscs exhibit a lyotropic liquid crystal behavior, in which the liquid crystalline phase depends on the temperature and concertation, and are suitable for RDC measurements.^22^

To demonstrate the feasibility of using nanodiscs as an alignment medium for NMR structural studies, uniformly ^15^N labeled cyt c was used as a model protein in this study. cyt c is a positively charged, water-soluble protein with pI of 9.6.^28^ We chose positively charged SMA-QA macro-nanodiscs to avoid any charge-charge interaction with cyt c.^29^ ^15^N labeled cyt c was added to a solution of macro-nanodiscs (40 mg/mL) and used in ^15^N-^1^H In-Phase, Anti-Phase Heteronuclear Single-Quantum Coherence (IPAP-HSQC)^30^ experiments to measure RDCs. 2D IPAP-HSQC spectra acquired at 20 °C showed the ^1^J_NH_-couplings measured in the presence and absence of nanodisc are comparable (Supporting information Figure S4), suggesting that the nanodiscs medium is isotropic at 20 °C. This is also confirmed from the appearance of an isotropic ^2^H peak for the same sample (Supporting information Figure S5). An anisotropic nanodiscs medium was obtained by increasing the temperature to 32 °C, for which the alignment was confirmed from the appearance of a doublet in deuterium NMR spectra (Figure 2A). Selected regions of 2D IPAP-HSQC spectra are shown in Figure 2C. RDCs were obtained by subtracting coupling (^1^J_NH_ or ^1^J_NH_ + D_NH_) values measured from 2D IPAP-HSQC spectra acquired at isotropic (20 °C) and anisotropic (32 °C) conditions. Our experimental results show that the RDC values can be scaled by simply increasing the nanodiscs concertation or the temperature as evident from the increased RDCs obtained at higher concentration: (±6 Hz for 40 mg/mL and ±40 Hz for 100 mg/mL) (see Figures 2(C and D), 3(A and B) and S6).

**Figure 2.**
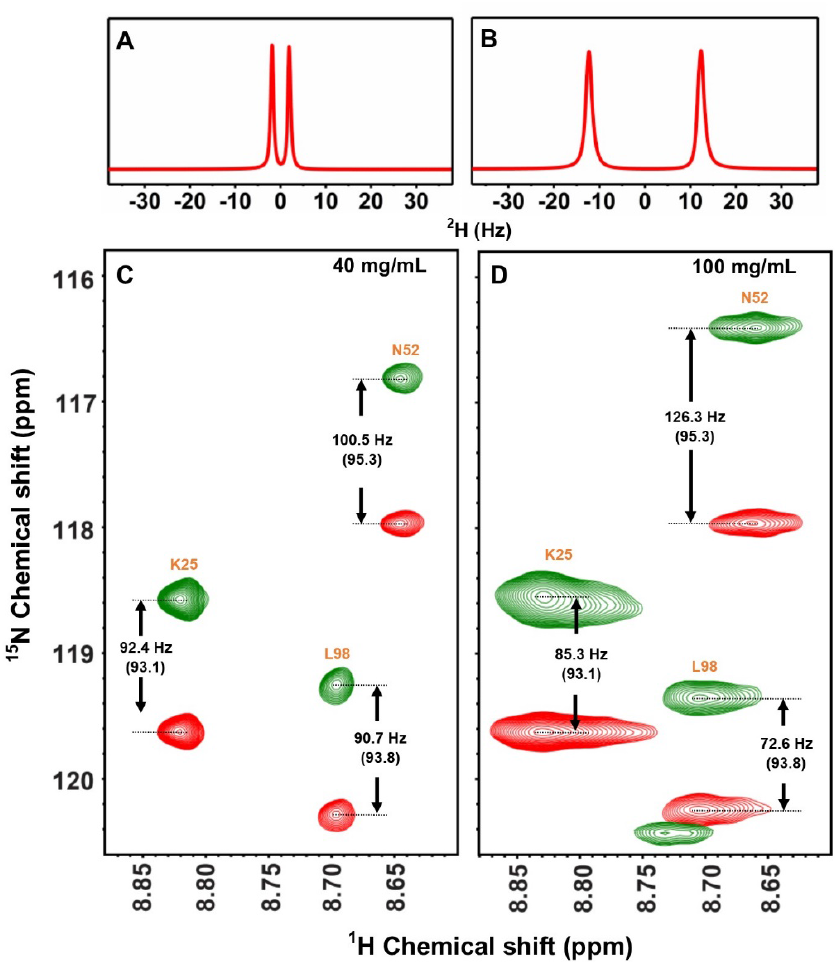
Deuterium NMR spectra of SMA-QA:DMPC macronanodiscs containing 200 μM ^15^N-cyt*C* in 10% D_2_O and phosphate buffer at 32 °C with a lipid concentration of 40 mg/mL (A) and 100 mg/mL (B). Selected region of 2D ^1^H-^15^N IPAP-HSQC spectra obtained using a lipid concentration of (C) 40 mg/mL and (D) 100 mg/mL at 32 °C in a 800 MHz NMR spectrometer. Isotropic ^1^J_NH_ values were obtained at 20 °C are shown in parenthesis (See supporting information Figure S5).

**Figure 3.**
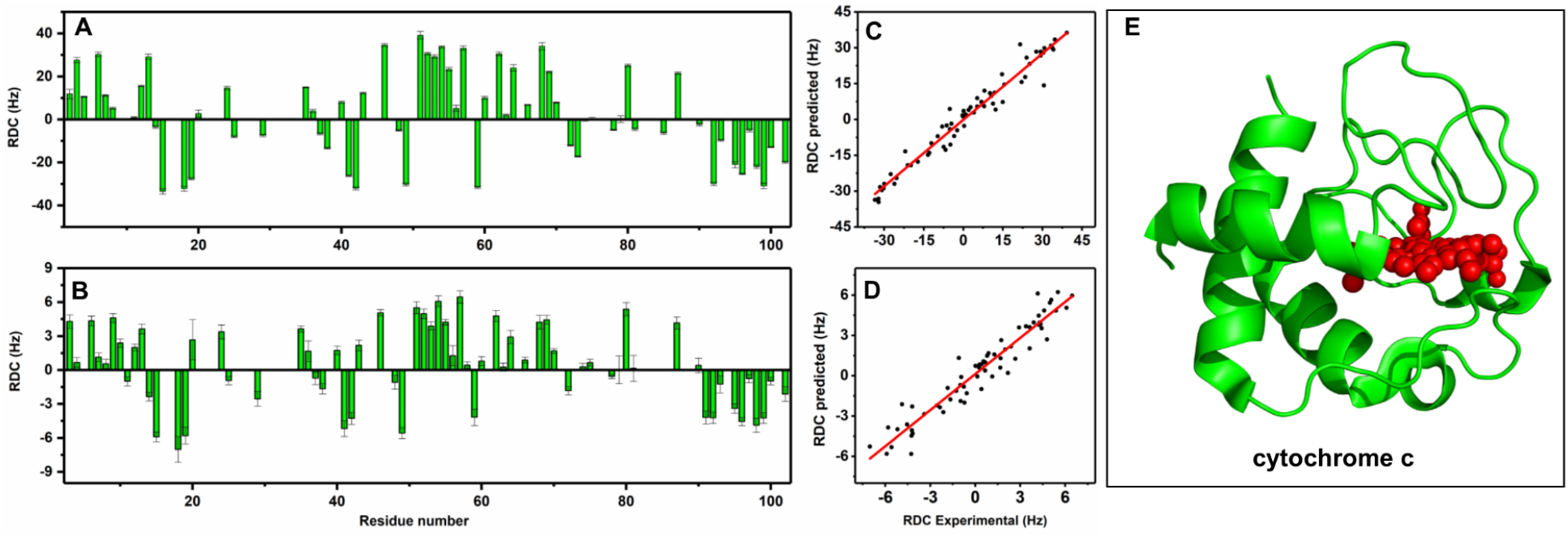
A) Experimentally measured residual dipolar couplings for each residue of ^15^N cyt c for lipid concentration of 100 mg/mL (A) and 40 mg/mL (B). Correlation of experimentally measured and calculated RDCs at 100 mg/mL (C) and 40 mg/mL (D). X-ray crystal structure of cyt c (PDB:6FF5).

To validate the accuracy of experimentally measured RDC values, the experimental RDCs were compared with RDCs calculated using reported crystal structures available of cyt c. Figure 3(C and D) shows the correlation between experimentally observed RDCs and calculated RDCs with R values 0.958 and 0.979 for 40 mg/mL and 100 mg/mL respectively. We further compared the experimental RDCs with those values calculated from different high-resolution structures reported for cyt c (Table S1, Figures S7 and S8). These good correlations further confirm that the macro-nanodiscs are useful for the measurement of RDCs.

In summary, we have successfully demonstrated the use of polymer macro-nanodiscs as an effective alignment medium for the measurement of RDC values in the structural studies using NMR experiments. Our results showed that the degree of alignment can be scaled by varying either the lipid-nanodiscs concentration or temperature. Using cyt c as a model system, in this study we showed that the experimentally measured RDCs are in good agreement with the calculated values from known structures of the protein. The pH resistant and divalent metal ion tolerant nature of SMA-QA nanodiscs can be further exploited to study a variety of proteins and other biomolecules for RDCs based NMR structural studies. The alignment tensor of the macro-nanodiscs alignment medium can easily be modulated by altering the interaction between the protein and the alignment medium by changing the lipid composition, and therefore it is feasible to measure RDCs with orthogonal tensors. Another advantage of polymer nanodiscs is that the protein (or any biomolecule) under study can be recovered from the sample using SEC after NMR experiments (See Figure S9). Due to these unique properties, we believe that the use of polymer nanodiscs as an alignment medium will be valuable for high-resolution structural studies on water-soluble biomolecules as well as to probe the interaction of molecules (such as peptides, proteins, ligands, RNA) with lipid membrane.

## ASSOCIATED CONTENT

### Supporting Information

A detailed description of materials and methods used in this study, and NMR spectra are included in the supporting information.

## Notes

The authors declare no competing financial interests.

## ACKNOWLEDGMENT

This study was supported by the funds from NIH (GM084018 to A.R.). We thank Dr. Nagao for help with the production of cytochrome c.

---

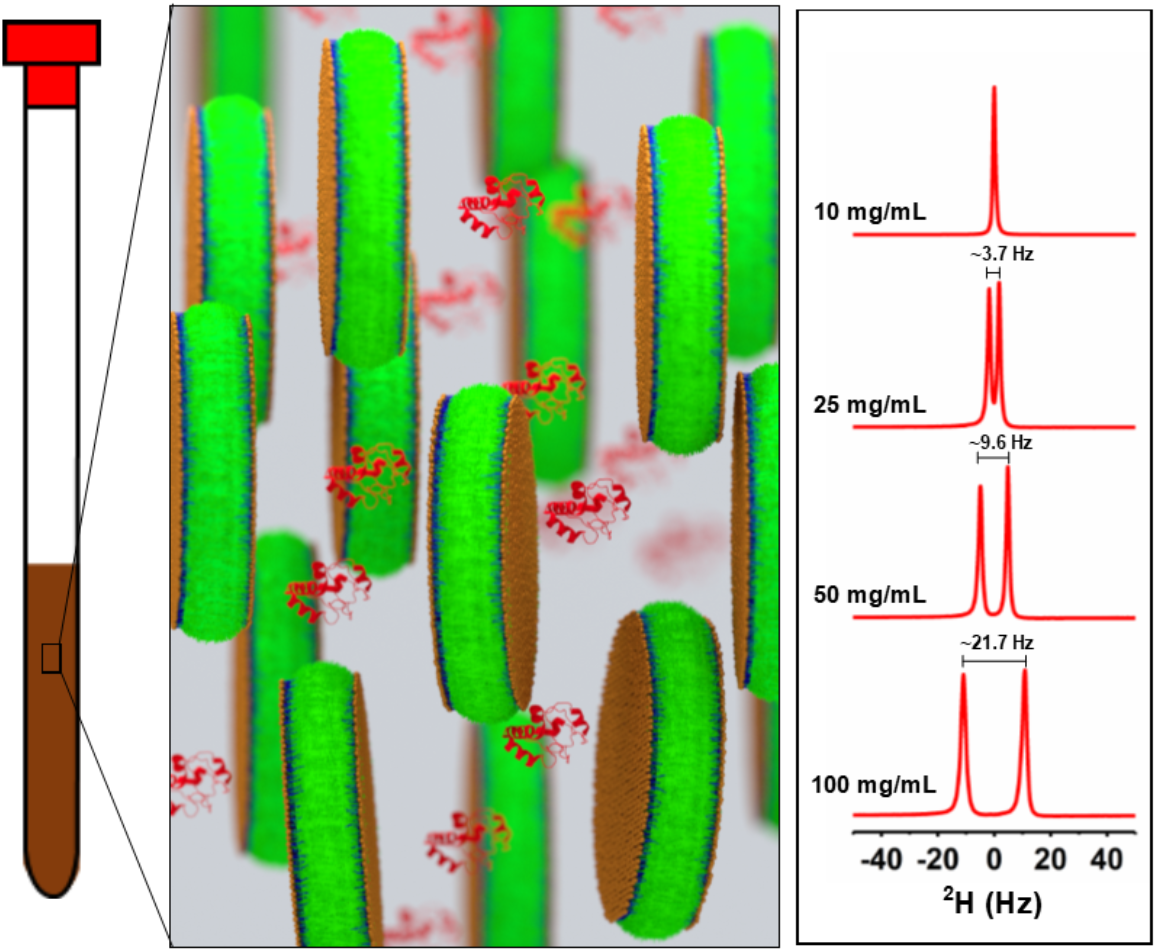
Table of Contents.

