## Supporting information for "Magnetic-alignment of polymer macro-nanodiscs enable residual dipolar couplings based high-resolution structural studies by NMR"

**Materials and Methods**

Poly(Styrene-co-Maleic Anhydride) cumene terminated (SMA), with a ~1.3:1 molar ratio of styrene:maleic anhydride and average molecular weight of $M_n$~1600 g/mol, N-Methyl-2-Pyrrolidone (NMP), 2-Aminoethanol (EA), Triethylamine ($Et_3N$), HEPES, potassium phosphate, acetic acid (HOAc), hydrochloric acid (HCl), and sodium hydroxide (NaOH) were purchased from Sigma-Aldrich®. 1,2-dimyristoyl-*sn*-glycero-3-phosphocholine (DMPC) was purchased from Avanti Lipids Polar, Inc®. Uniformly $^{15}$N-labeled cyt *c* was expressed and purified as reported earlier the literature.[1]

**Synthesis of SMA-QA:** SMA-QA was synthesized as described previously.[2] Briefly, 1 g of SMA and 1.3 g of (2-aminoethyl)trimethyl ammonium chloride hydrochloride was added to anhydrous DMF (30 mL), followed by the addition of triethylamine (5 mL). The reaction mixture was stirred at 80 °C for 12 h. The reaction mixture is cooled to room temperature and the product was precipitated and washed 3 times using diethyl ether. The resulting product was dried under vacuum to obtain a white powder. The resulting powder was added to 30 mL of acetic anhydride and followed by the addition of 660 mg of sodium acetate and 0.275 mL of triethylamine. The reaction mixture was stirred at 80 °C for 12 h. The reaction mixture was cooled to room temperature, precipitated, and washed three times with diethyl ether and dried under vacuum. The product was dissolved in water, passed through the Sephadex LH-20 column. The product was collected and lyophilized to obtain a brown powder.

**Preparation of nanodiscs:** SMA-QA stock solutions were prepared by dissolving 100 mg of SMA-QA in 10 mL of 20 mM potassium phosphate 50 mM NaCl (pH 7.4). DMPC stock solutions were prepared by the addition of 100 mg of DMPC powder in 10 mL of 20mM potassium phosphate 50 mM NaCl (pH 7.4) and subjected to three freeze-thaw cycles to get a milky white solution. Nanodiscs were prepared by the addition of polymer to DMPC solution with weight ratios corresponding to 1:0.5 w/w of DMPC and SMA-QA, and incubated overnight at 30 °C. The resulting nanodiscs were subjected to Size Exclusion Chromatography to separate the nanodiscs from the free polymer. The nanodiscs fractions were collected (Figure S1), and 10 kDa Amicon® filters were used to make a required lipid concentration. In all experiments, concertation was estimated based on the starting lipid concentration. Nanodiscs sizes were measured using DLS and TEM.

**Size-Exclusion Chromatography (SEC):** Manually packed Superdex® 200 16/600 GL semi preparative column attached to an AKTA® purifier Fast Protein Liquid Chromatography (FPLC) purification system (GE Healthcare®) with a 5 mL loop was used to elute the initial nanodisc samples with a 20 mM potassium phosphate and 50 mM NaCl buffer (pH 7.4) at a flow rate of 1.5 mL/min. Detection was done by collecting the absorbance at λ = 254 nm. cyt c separation from the nanodiscs was performed using Superdex® 200 10/600 GL column. 1 mL loop was used for injecting the sample, 1.2 mL/min flow rate was used for the elution, and the detection was done by using absorbance at λ = 280 nm.

**RDC measurements using In-Phase, Anti-Phase Heteronuclear Single-Quantum Coherence (IPAP-HSQC):** NMR samples were prepared by the addition of $^{15}$N- cyt c to the solution containing macro-nanodiscs to give a final concentration of 200 μM protein and 40 mg/mL or 100 mg/mL lipid concentration; 10% $D_2O$ was added to all NMR samples for locking. Deuterium NMR spectra were acquired to check the degree of magnetic-alignment of macro-nanodiscs. Two-dimensional spectra $^{15}$N/$^{1}$H IPAP-HSQC were acquired on a Bruker 800 MHz NMR spectrometer equipped with a 5 mm triple resonance TCI cryoprobe. Spectra were acquired using the following parameters: number of scans 4; 2048 t2*512 t1 points (for each IP and AP); 1.2 s recycle delay. The spectra were split into two 2D spectra using *split ipap2* AU macro with a default setting in topspin. The resulting spectra were processed using Topspin with zero filling up to 4028*4028 points. Peaks were assigned based on the previous assignments.[3] Peak position and Line widths (LW) were obtained using the integration module with a default setting in Sparky for each peak. Peaks with signal-to-noise ratio (S/N) above 10 were used for further analysis. RDCs were calculated by subtracting the $^{1}J_{HN}$(anisotropic)-$^{1}J_{HN}$ (isotropic) using Excel. Lower limit for the accuracy in the measurement of a peak position is calculated by the ratio of LW/(S/N).[4] cyt c structural coordinates were obtained from the RCSB protein data bank,[5] hydrogens were added using the web server[6]. Alignment tensors were obtained using MODULE software.[7] Correlations between the measured and calculated RDCs are represented by Pearson correlation coefficient (R) obtained using the OriginLab© software.[8]

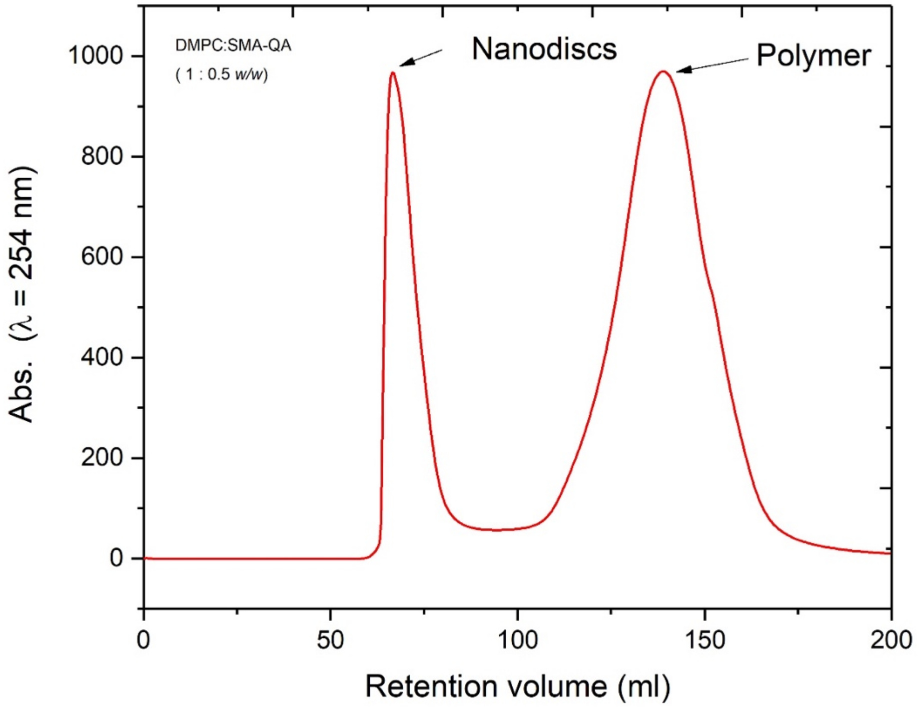


**Figure S1:** Size exclusion chromatogram of 1:0.5 w/w DMPC:SMA-QA. Nanodisc and polymer fractions are indicated by arrows.

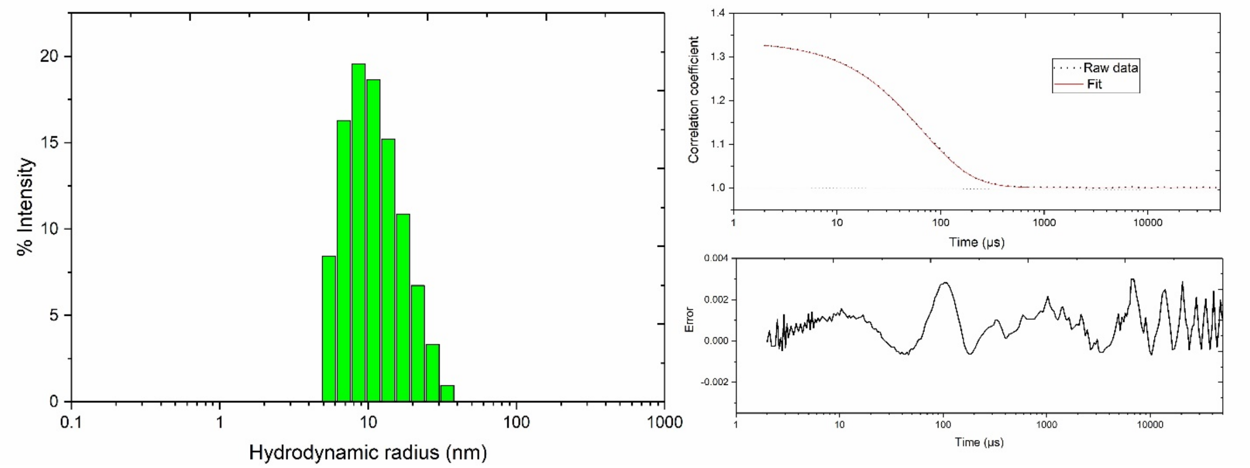


**Figure S2:** (Left) Dynamic Light Scattering spectra of nanodiscs (1:0.5 w/w SMA-QA DMPC) after SEC. (Right) Autocorrelation function used to estimate the hydrodynamic radius and their fits are shown; residuals of the fitting are shown at the bottom.

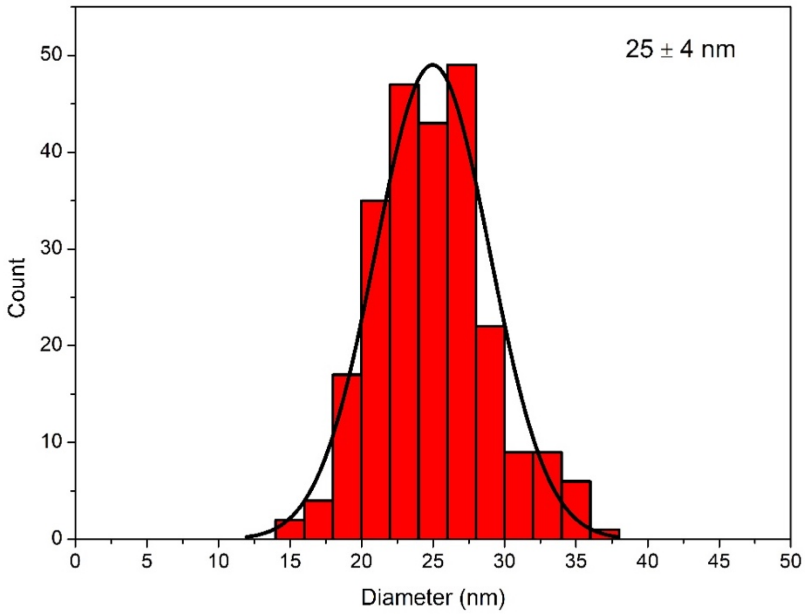


**Figure S3:** Size distribution and Gaussian fitting of nanodiscs calculated from TEM images. Sizes were measured using ImageJ[9] software by manually picking the particles.

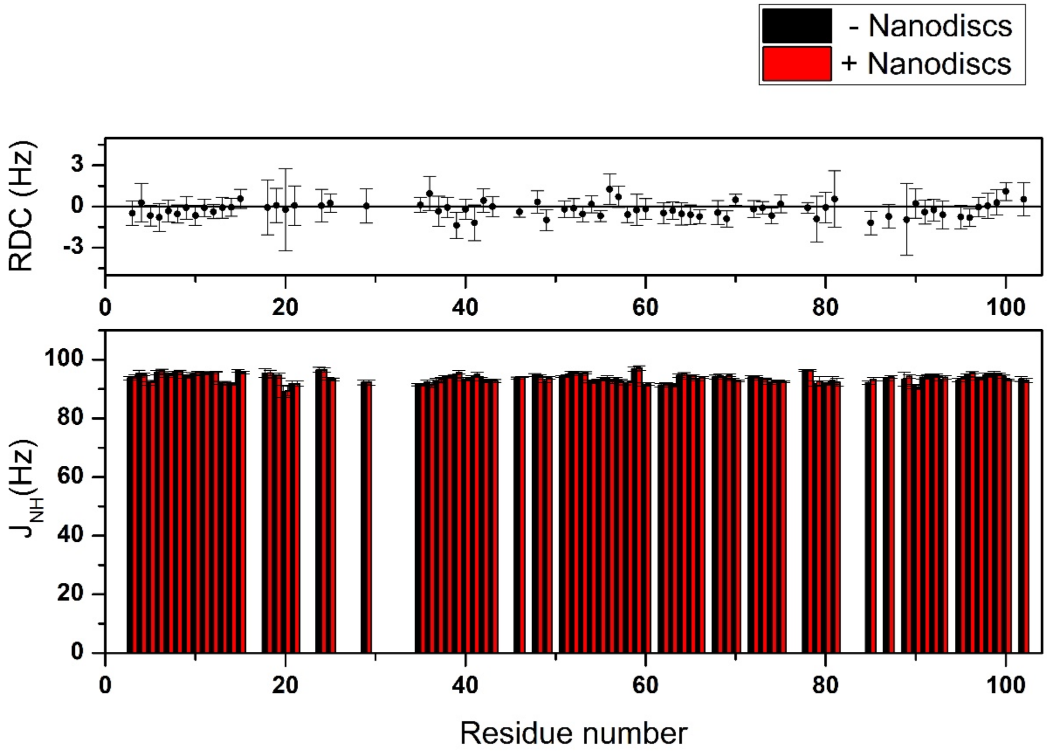


**Figure S4:** $^1J_{NH}$ measured from IPAP-HSQC NMR spectra of $^{15}$N-cyt c (200 μM) in buffer (20 mM KPi, 50 mM NaCl, pH 7.4) at 25 °C and in the presence of DMPC-SMAQA nanodiscs (40 mg/mL) at 20 °C. Error bars represent the ratio of linewidth and signal to noise ratio. Difference between the couplings measured under these two conditions are plotted on top. The $^1J_{NH}$ couplings measured in the absence of nanodiscs are comparable to those obtained in the presence of the nanodiscs, showing the isotropic nature of nanodiscs samples at 20 °C.

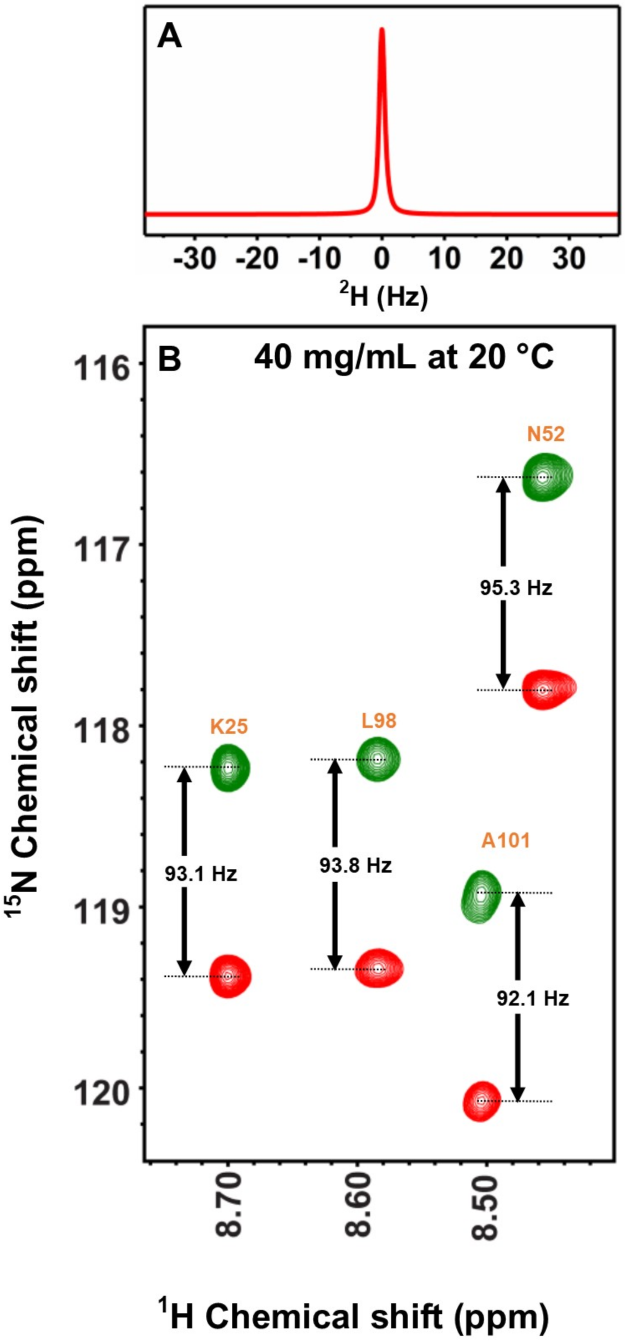


**Figure S5:** (A) Deuterium NMR spectrum of nanodiscs containing 200 µM $^{15}$N-cyt c in 10% $D_2O$ and in phosphate buffer containing 40 mg/mL lipid concentration of macro-nanodiscs at 20 °C. (B) Selected region (for the residues shown in Figure 2 in the main text) $^1$H-$^{15}$N IPAP-HSQC NMR spectra of 200 µM $^{15}$N-cyt c in SMA-QA:DMPC macro-nanodiscs at 20 °C.

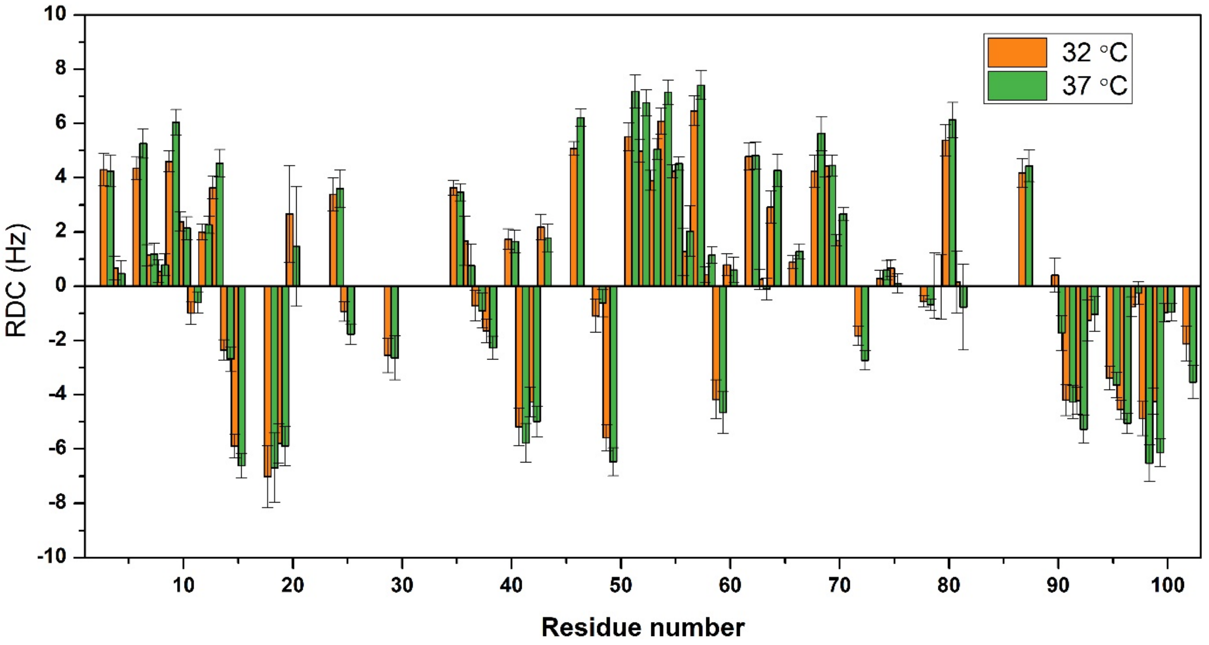


**Figure S6:** Residual dipolar couplings measured from $^{15}$N-cyt c at two different temperatures as indicated. 40 mg/mL of lipid concentration of DMPC-SMAQA macro-nanodiscs was used. The RDCs measured at 37 °C are higher than those values obtained at 32 °C showing that the RDCs can be linearly scaled by varying the temperature of the sample.

| S.NO | PDB ID | Title | Reference | Resolution | Correlation *R* | Technique used |
|---|---|---|---|---|---|---|
| **1** | **1HRC** | High-resolution three-dimensional structure of horse heart cytochrome c | Bushnell, G.W., Louie, G.V., Brayer, G.D.<br><br>(1990) J.Mol.Biol. 214: 585-595 | 1.9 Å | 0.958 | X-ray |
| **2** | **6FF5*** | X-ray structure of bovine heart cytochrome c at high ionic strength | Merlino, A. (2018) Biometals 31: 277-284 | 1.74 | 0.979 | X-ray |
| **3** | **1CRC** | Cytochrome c at low ionic strength | Sanishvili, R., Volz, K.W., Westbrook, E.M., Margoliash, E. (1995) Structure 3: 707-716 | 2.08 | 0.917 | X-ray |
| **4** | **3O1Y** | Electron transfer complexes: Experimental mapping of the redox-dependent cytochrome c electrostatic surface | De March, M., Demitri, N., De Zorzi, R., Casini, A., Gabbiani, C., Guerri, A., Messori, L., Geremia, S. (2014) J. Inorg. Biochem. 135: 58-67 | 1.75 | 0.969 | X-ray |
| **5** | **1AKK** | Solution structure of oxidized horse heart cytochrome c, NMR, minimized average structure | Banci, L., Bertini, I., Gray, H.B., Luchinat, C., Reddig, T., Rosato, A., Turano, P. (1997) Biochemistry 36: 9867-9877 | N/A | ND | NMR |
| **6** | **5C0Z** | The structure of oxidized rat cytochrome c at 1.13 angstroms resolution | Edwards, B.F.P., Mahapatra, G., Vaishnav, A.A., Brunzelle, J.S., Huttemann, M. (to be published) | 1.13 | 0.973 | X-ray |

* Sequence similarity 98%

**Table S1:** Details of cyt c structures used for the comparison of the calculated and experimentally measured RDC values. All the crystal structures exhibit very good correlation (with R>0.9) except for 1AKK. Correlation plots are shown in Figure S7.

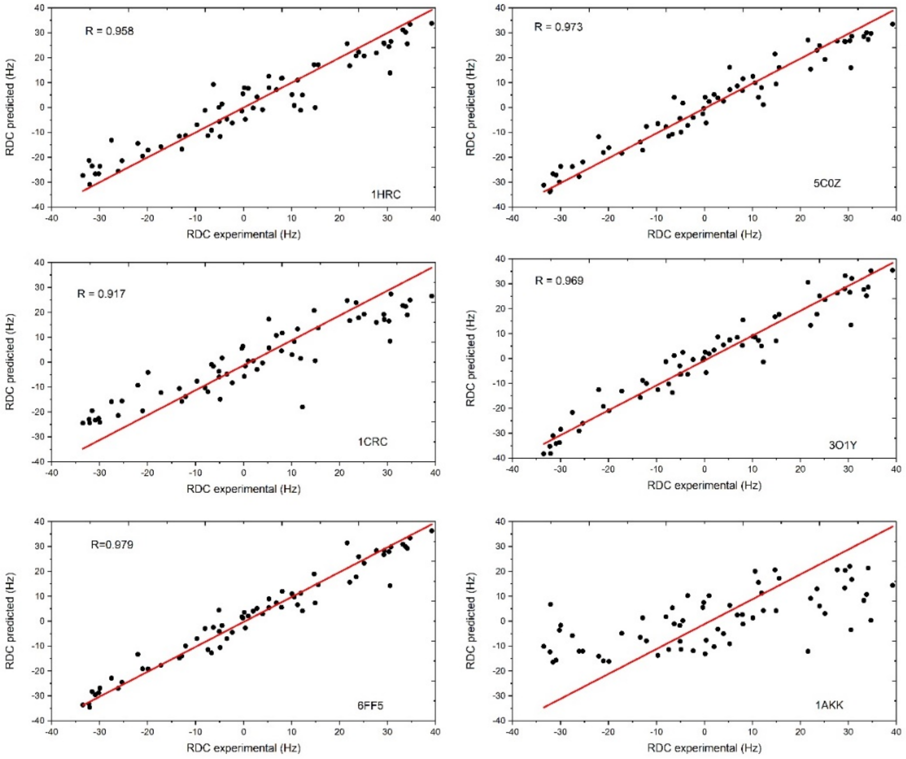


**Figure S7**: Correlation of experimentally measured and calculated RDCs for $^{15}$N-cyt c (200 μM) in macro-nanodiscs (100 mg/mL of lipid concentration) for the structures mentioned in Table S1.

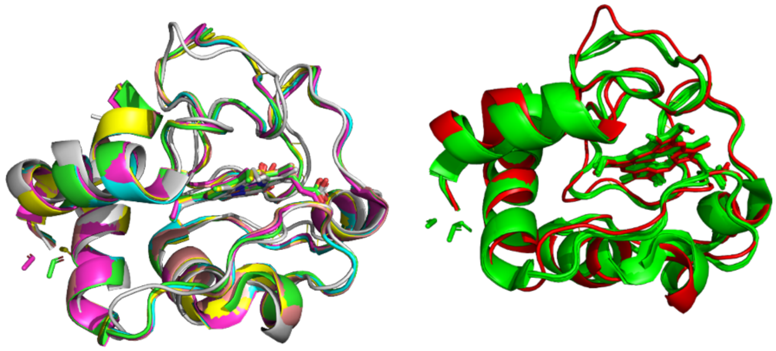

**Figure S8:** Cartoon representation of cyt c structures used for the correlation in Figure S7 (left). The backbone RMSD is 1.03 Angstroms. The structural variations in PDB ID 1AKK (red) compared to other crystal structures (green) are shown in right. Structures were aligned, and RMSD was calculated using PyMOL software.[10]

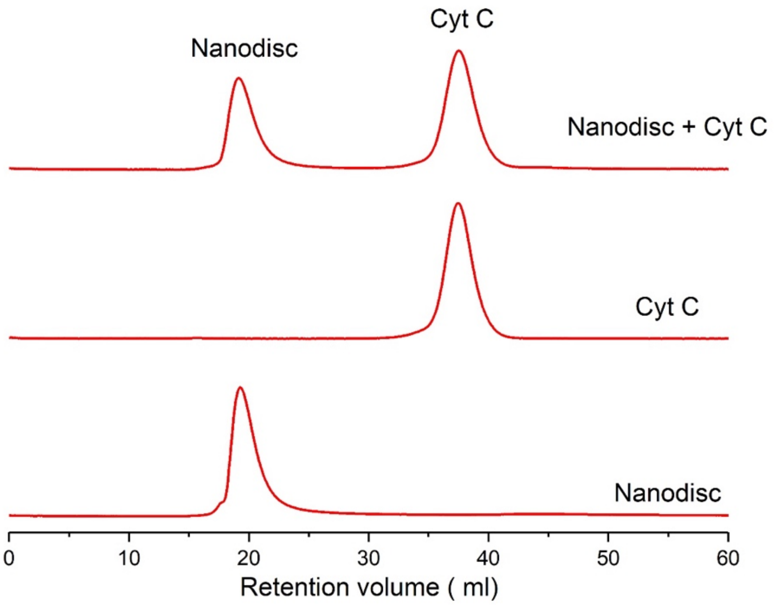


**Figure S9:** SEC profiles showing the elution volume differences between macro-nanodiscs (20 ml) and cyt c (40 mL). This suggest that the reconstituted protein can be recovered after the measurement of RDC values. Detection was done using absorbance at λ = 280 nm.

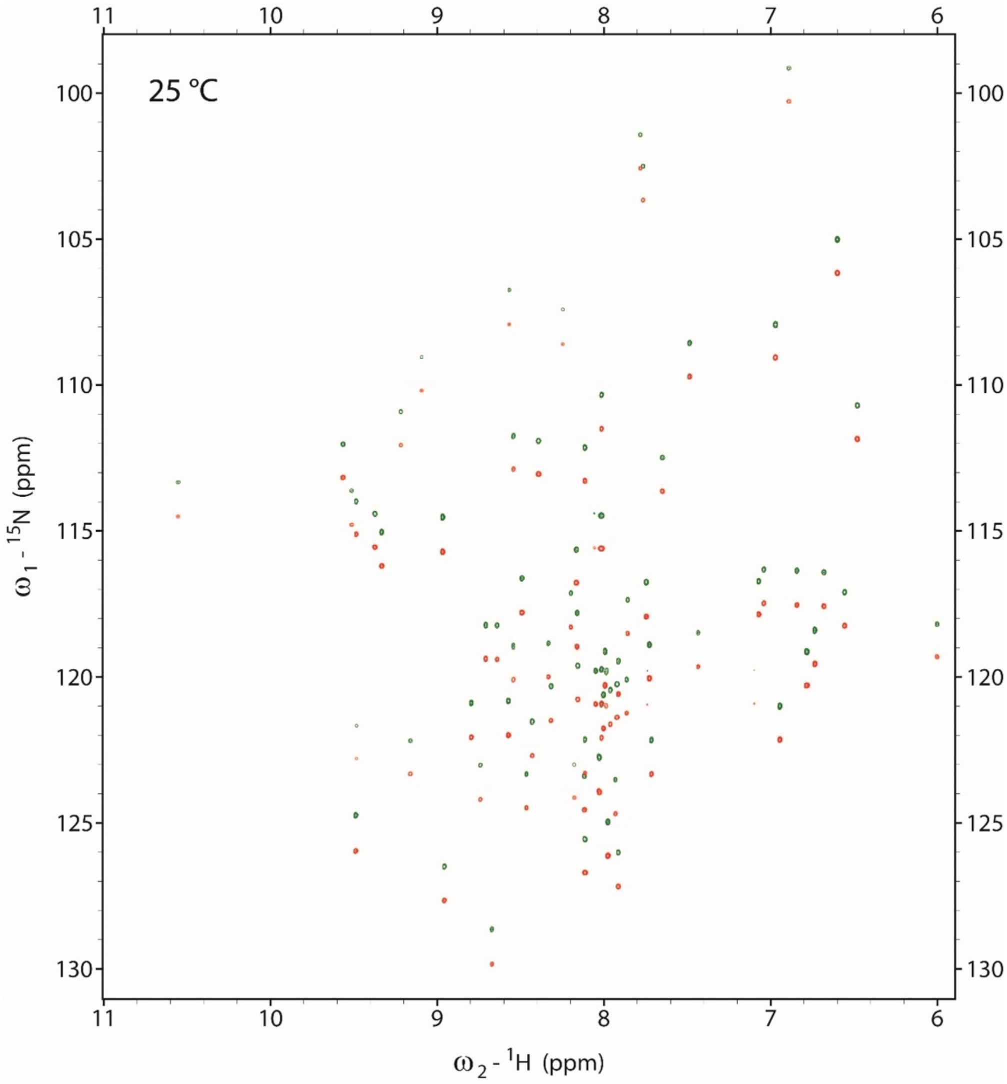


**Figure S10:** 2D IPAP-HSQC NMR spectra of $^{15}$N-cyt c (200 μM) in 10% $D_2O$, 20 mM Kpi, 50 mM NaCl pH 7.4 obtained at 25 °C.

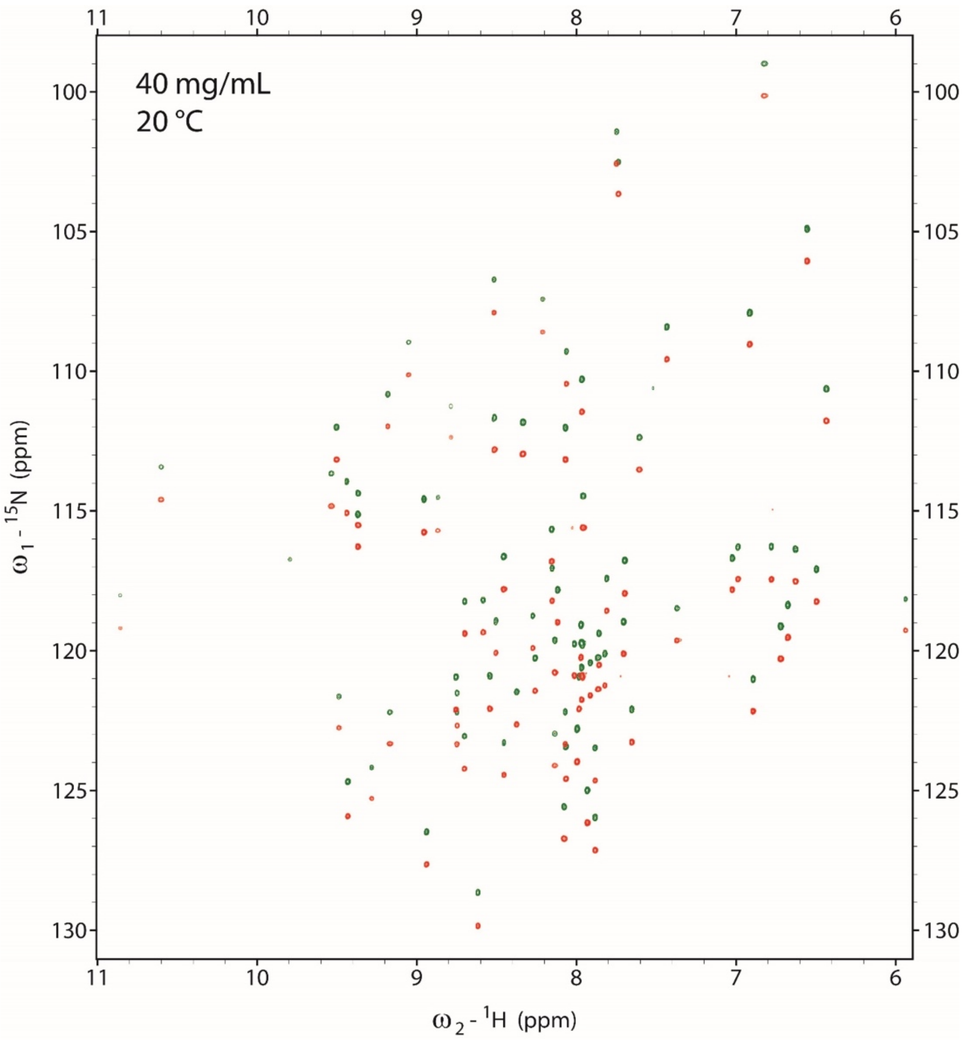


**Figure S11:** 2D IPAP-HSQC NMR spectra of [15]N-cyt c (200 μM) in SMA-QA:DMPC macro-nanodiscs (40 mg/mL) in 10% $D_2O$, 20 mM Kpi, 50 mM NaCl pH 7.4 obtained at 20 °C.

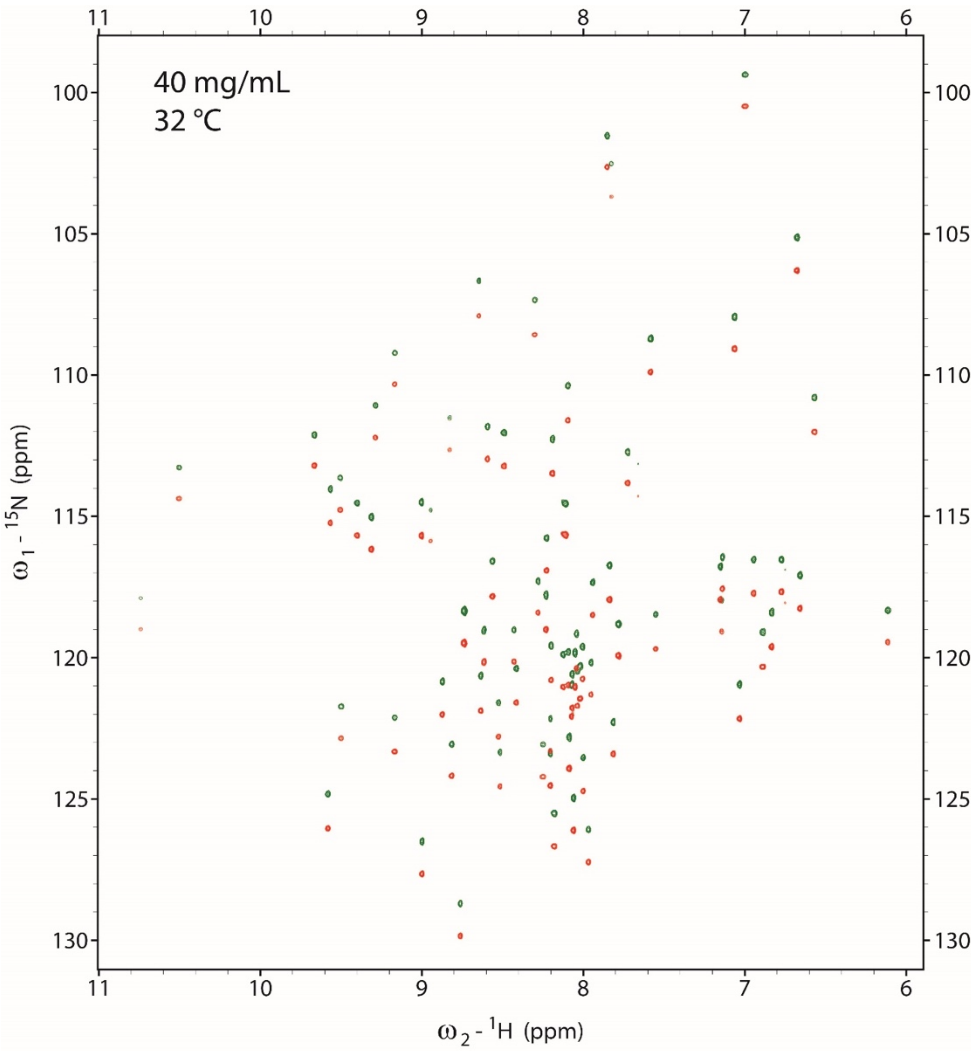


**Figure S12:** 2D IPAP-HSQC NMR spectra of $^{15}$N-cyt c (200 μM) in SMA-QA:DMPC macro-nanodiscs (40 mg/mL) in 10% $D_2O$, 20 mM Kpi, 50 mM NaCl pH 7.4 obtained at 32 °C.

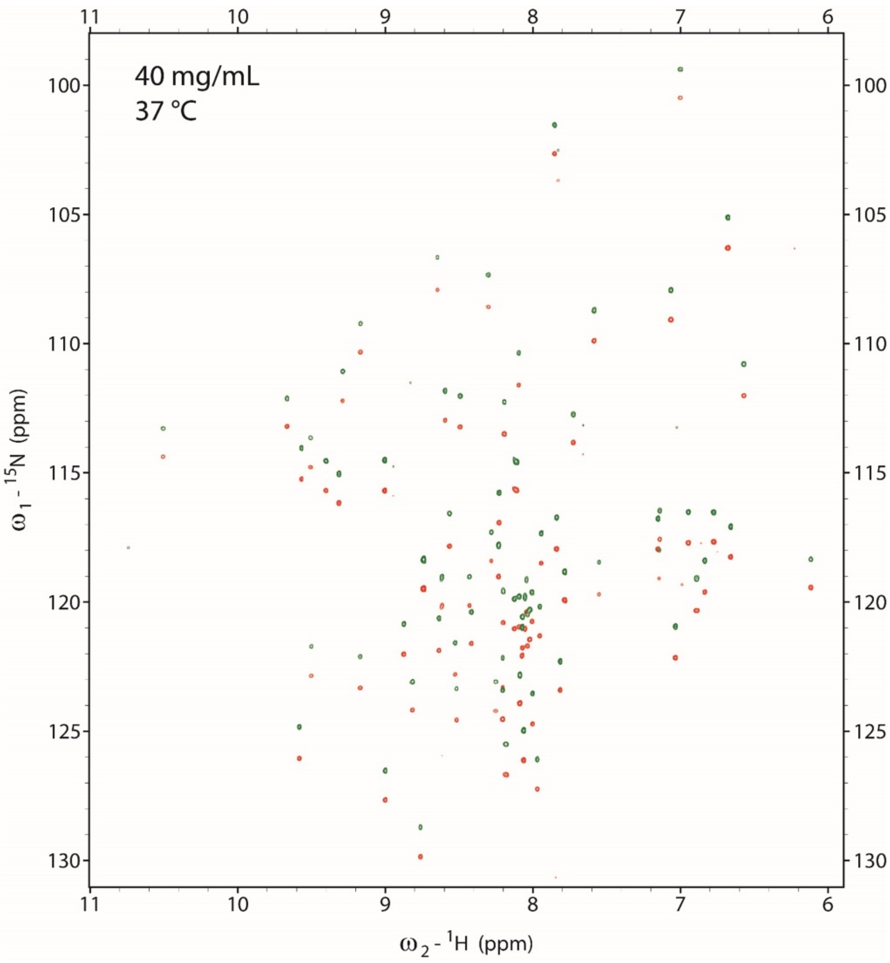


**Figure S13:** 2D IPAP-HSQC NMR spectra of $^{15}$N-cyt c (200 μM) in SMA-QA:DMPC macro-nanodiscs (40 mg/mL) in 10% $D_2O$, 20 mM Kpi, 50 mM NaCl pH 7.4 obtained at 37 °C.

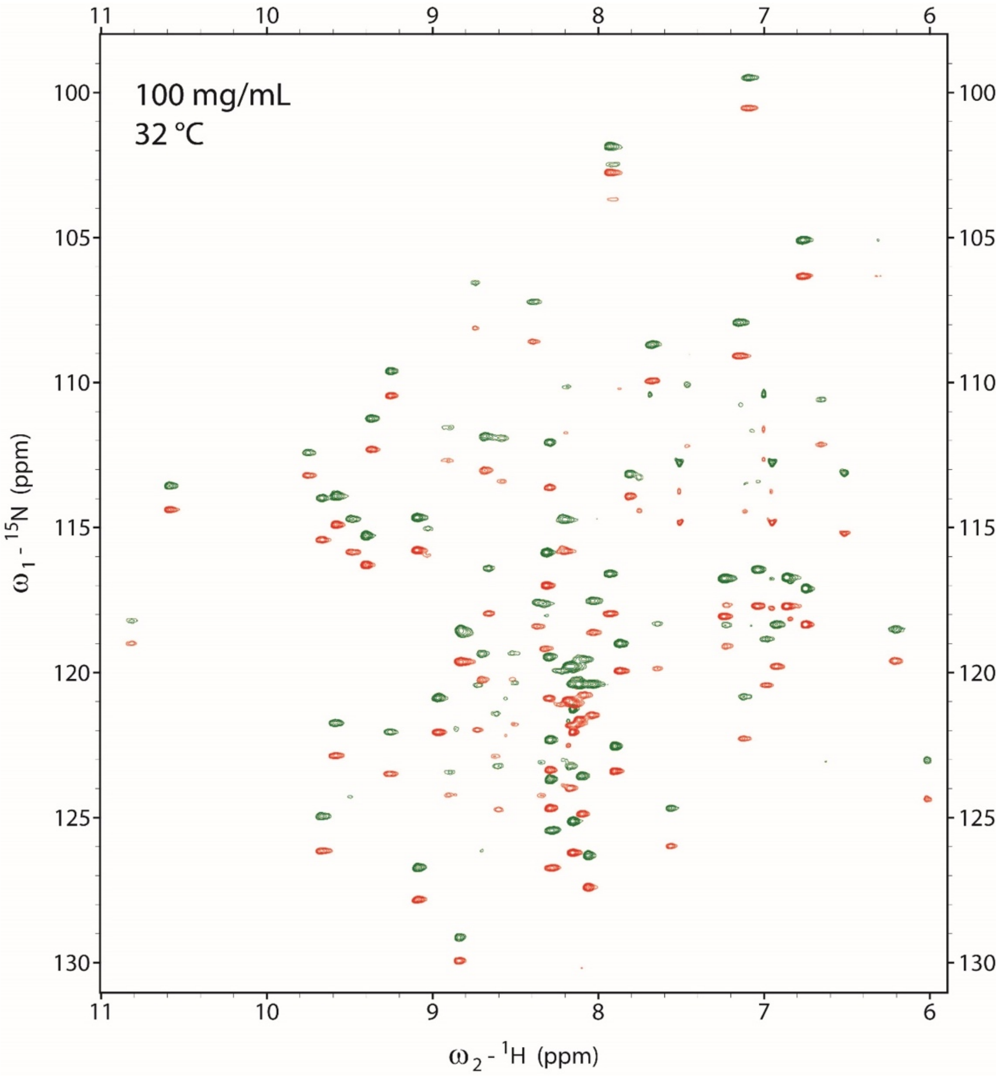


**Figure S14:** 2D IPAP-HSQC NMR spectra of $^{15}$N-cyt c (200 μM) in SMA-QA:DMPC macro-nanodiscs (100 mg/mL) in 10% $D_2O$, 20 mM Kpi, 50 mM NaCl pH 7.4 obtained at 32 °C.
